# Osmotic and ionic regulation by hololimnetic freshwater crustaceans: a molecular role for gill ion transporters

**DOI:** 10.1101/2020.04.17.046698

**Authors:** Milene Mantovani, John Campbell McNamara

## Abstract

Owing to their extraordinary niche diversity, the Crustacea are ideal for comprehending the evolution of osmoregulation. The processes that effect systemic hydro-electrolytic homeostasis maintain hemolymph ionic composition via membrane transporters located in highly specialized gill ionocytes. We evaluated physiological and molecular hyper- and hypo-osmoregulatory mechanisms in two phylogenetically distant, freshwater crustaceans, the crab *Dilocarcinus pagei* and the shrimp *Macrobrachium jelskii*, when osmotically challenged for up to 10 days. When in distilled water, hemolymph osmolality and [Cl^−^] increased briefly in *D. pagei*, stabilizing at initial values, while [Na^+^] decreased continually. Gill V(H^+^)-ATPase, Na^+^/K^+^-ATPase and Na^+^/K^+^/2Cl^−^ gene expressions were unchanged. In *M. jelskii*, hemolymph osmolality, [Cl^−^] and [Na^+^] decreased continually for 12 h, the shrimps no longer surviving. Gill transporter gene expressions increased 2- to 5-fold. After 10-days exposure to brackish water, *D. pagei* was isosmotic, iso-chloremic and iso-natriuremic. Gill V(H^+^)-ATPase expression decreased while Na^+^/K^+^-ATPase and Na^+^/K^+^/2Cl^−^ expressions were unchanged. In *M. jelskii*, the hemolymph was hypo-regulated, particularly [Cl^−^]. Transporter expressions initially increased 3- to 12-fold, declining to control values. Gill V(H^+^)-ATPase expression underlies the ability of *D. pagei* to survive in fresh water while Na^+^/K^+^-ATPase and Na^+^/K^+^/2Cl^−^ expressions enable *M. jelskii* to deal with osmotic challenge. These findings reveal divergent responses in two unrelated crustaceans habiting a similar osmotic niche. While *D. pagei* has maintained the capacity to tolerate elevated cellular isosmoticity despite its inability to secrete salt, *M. jelskii* displays clear hypo-osmoregulatory ability. Each species has developed distinct strategies at the transcription and systemic levels during adaptation to fresh water.

**Summary statement:** During their evolutionary adaptation to fresh water, unrelated hololimnetic crustaceans have developed physiological strategies like tolerating elevated cellular isosmoticity or regulating hypo-osmoregulatory ability at the gene transcription level.

## Introduction

Water is a fundamental component of the intra- and extracellular fluids, and its osmotic movement between this internal *milieu* and the external environment, be it obligatory or regulated, can constitute a severe challenge to organisms (Péqueux, 1995; Schmidt-Nielsen, 2002; Willmer et al., 2005; McNamara and Faria, 2012). Hydro-electrolytic homeostasis is, therefore, a major physiological determinant of potential niche diversification.

The more inclusive clades of decapod crustaceans all derive from marine ancestors (Ruppert and Barnes, 1994; Freire et al., 2013). At different moments since the early Carboniferous some 350 million years ago, certain shrimp-like taxa began to invade brackish and fresh waters (Cohen, 2003). The trichodactylid, potamoid and grapsid crabs invaded between 70 and ≈87 Mya, and the caridean shrimps about ≈70 Mya (Tsang et al., 2014; Anger, 2013). Crustacean species that inhabit dilute media or fresh water (<0.5 g salt/L [salinity, ‰S]) are subject to intense osmotic water influx and passive salt efflux across their body surfaces, and to salt loss in their urine. Such loss is counterbalanced by powerful hyper-osmoregulatory mechanisms, such as active salt uptake (Onken and Putzenlechner, 1995; Weihrauch et al., 2004; Freire et al., 2008a), and by producing voluminous, often dilute urine. These capabilities, together with the a calcified exoskeleton, reduction in epithelial permeability (Kirschner, 1991; Péqueux, 1995; Charmantier et al., 2009; Ashelby et al., 2012) and tight regulation of the composition and volume of the intra- and extracellular fluids (Péqueux, 1995; Augusto et al., 2007a,b; Freire et al., 2008b) constitute vital morphological and physiological adaptations that likely have subsidized the occupation of dilute media (Mantel and Farmer, 1983; Péqueux, 1995). Hololimnetic crustaceans in particular are highly tolerant of this severe osmotic challenge and are excellent hyper-osmoregulators, establishing elevated osmotic and ionic gradients against fresh water (≈30: 1).

The main processes that effect osmotic and ionic homeostasis in freshwater Crustacea are: (i) Isosmotic Intracellular Regulation, that maintains the composition and volume of the intracellular fluid (Péqueux, 1995; Augusto et al., 2007b; Freire et al., 2013), mediated by the transport of ions such as Na^+^, K^+^ and Cl^−^ (Freire et al., 2013) and by the synthesis and/or degradation of amino acids and peptides, reducing or increasing intracellular osmolality; and (ii) Anisosmotic Extracellular Regulation, that holds the osmolality, ionic concentration and volume of the hemolymph within species-specific limits, through the action of ion transporting proteins like the Na^+^/K^+^- ATPase and V(H^+^)-ATPase (Towle and Kays, 1986; Tsai and Lin, 2007), and membrane transporters such as the Na^+^/K^+^/2Cl^−^ symporter, Na^+^/H^+^ antiporter and ion channels, located in specialized ionocytes in the gills, antennal glands and intestine (Péqueux, 1995; Freire et al., 2008a; McNamara et al., 2005; McNamara et al., 2015).

Crustacean gills are the principal sites of ion and gas exchange and pH regulation. Their ionocytes undertake active ion transport, and are characterized by an extensive surface area exhibiting a cell membrane highly amplified by apical evaginations, and abundant, deep basal invaginations, each closely associated with mitochondria (Freire and McNamara, 1995; McNamara and Lima, 1997; Onken and McNamara, 2002; Weihrauch et al., 2004; Furriel et al., 2010; McNamara and Faria, 2012; McNamara et al., 2015). The different osmoregulatory patterns seen in the Crustacea result from distinct transporter arrangements in the apical and basal ionocyte membranes (McNamara and Faria, 2012).

In strong freshwater hyper-osmoregulators (Péqueux, 1995; Kirschner, 2004; Freire et al., 2008a; McNamara and Faria 2012), carbonic anhydrase in the ionocyte cytosol generates H^+^ and HCO_3_^−^ from metabolic CO_2_ used by the apical V(H^+^)-ATPase and Cl^−^/HCO_3_^−^ antiporter. The apical membrane, hyperpolarized by H^+^ extrusion, allows Na^+^ influx via apical Na^+^ channels while the Na^+^/K^+^-ATPase located in the basal membrane (Towle and Kays, 1986) actively transports Na^+^ to the hemolymph in exchange for K^+^. HCO_3_^−^ is exchanged for Cl^−^ via the apical Cl^−^/HCO_3_^−^ antiporter, Cl^−^ flowing through basal Cl^−^ channels to the hemolymph (Putzenlechner et al., 1992; Genovese et al., 2005).

Some species exhibit hypo-osmoregulatory ability (Mantel and Farmer, 1983; Péqueux, 1995; Freire et al., 2003) in which salt secretion is driven by the basally located Na^+^/K^+^-ATPase and Na^+^/K^+^/2Cl^−^ symporter. Hemolymph Na^+^, K^+^ and Cl^−^ are transported down the Na^+^ gradient into the cytoplasm through the symporter. Na^+^ returns via the Na^+^/K^+^-ATPase while K^+^ recycles through basal K^+^ channels, creating a negative cell potential that drives Cl^−^ efflux through apical Cl^−^ channels. The negative transepithelial voltage generated in the subcuticular space drives paracellular Na^+^ efflux (McNamara and Faria, 2012).

The investigation of ion transport by gill ionocytes and specific ion transporters like the Na^+^/K^+^-ATPase, V(H^+^)-ATPase, Na^+^/K^+^/2Cl^−^ symporter, Na^+^/H^+^ antiporter and carbonic anhydrase, in response to osmotic challenge, has focused recent osmoregulatory research on functional gene expression studies (Faleiros et al., 2010; Moshtaghi et al., 2018; Rahi et al., 2017). These transporters are particularly abundant in gill ionocytes (McNamara and Torres, 1999; Boudour-Boucheker et al., 2014; Maraschi et al., 2015; Faleiros et al., 2017, 2018), and adjustments at the molecular level in enzyme activity, in gene and protein expressions and in localization underlie systemic physiological responses to the osmolality of the external medium (Lima et al., 1997; Weihrauch et al., 2004; Santos et al., 2007; Belli et al., 2009; Faleiros et al., 2010; Boudour-Boucheker et al., 2014; França et al., 2013; Havird et al., 2013; Maraschi et al., 2015; Faleiros et al., 2017; Freire et al., 2018).

In the present investigation, we examine systemic and molecular osmoregulatory adjustments using two phylogenetically distant freshwater decapod crustaceans as models: the brachyuran crab *Dilocarcinus pagei* (Stimpson, 1861) and the caridean shrimp *Macrobrachium jelskii* (Miers, 1877). Both are excellent hyper- osmoregulators and do not depend on brackish water to complete their life cycles (Magalhães, 2000; Onken and McNamara, 2002), characterizing them as hololimnetic species. Anisosmotic extracellular regulation is the predominant osmoregulatory mechanism in these species.

*Dilocarcinus pagei* is a trichodactylid brachyuran, endemic to the Amazon, Paraguay and Paraná river basins of South America. This crab is an ancient and well- adapted inhabitant of fresh water, exhibiting strong anisosmotic and anisoionic hyper- regulatory ability (Onken and McNamara, 2002; Augusto et al., 2007b) with low isosmotic and isoionic points, although it does not produce dilute urine (Augusto et al., 2007b). The gills are phyllobranchiate, the posterior gill lamellae being structurally and functionally differentiated from the anterior lamellae (Onken and McNamara, 2002). Morphological and electrophysiological findings, and semi- quantitative gene expression of ion transporters confirm that the three posterior gills are the main sites of ion uptake (Onken and McNamara, 2002; Weihrauch et al., 2004). In *D. pagei*, the asymmetrical, opposing epithelia of the posterior gill lamellae are separated by an irregular hemolymph space traversed by pillar cell pericarya. The thin distal epithelium consists of well developed apical flanges projecting from the pillar cell pericarya. The thick proximal epithelium consists of cuboid ionocytes, exhibiting extensive basal invasions (Weihrauch et al., 2004; Furriel et al., 2010). *Dilocarcinus pagei* exhibits active and independent Na^+^ and Cl^−^ uptake, the posterior gill epithelium being responsible for maintaining elevated osmotic and ionic gradients (≈30: 1) against fresh water. Active Na^+^ transport is driven across the thick proximal epithelium by the Na^+^/K^+^-ATPase, while Cl^−^ transport is thought to take place across the thin distal epithelium, driven by V(H^+^)-ATPase activity (Onken and McNamara, 2002; Weihrauch et al., 2004).

The caridean shrimp *Macrobrachium jelskii* is widely endemic throughout South America and is distributed within most large Brazilian river basins, like the Amazon, Tocantins/Araguaia, São Francisco and Paraná/Paraguay watersheds (Magalhães et al., 2005; Boos et al., 2012; Pileggi et al., 2013). The palaemonid genus Macrobrachium includes many freshwater shrimp species distributed throughout tropical and subtropical zones (Holthuis, 1980). Diadromous Macrobrachium species inhabit fresh water as adults but depend on salt water for larval development; hololimnetic species are restricted entirely to fresh water. All are strong hyper- osmoregulators (Denne, 1968; Moreira et al., 1983; Charmantier and Anger, 1999; Freire et al., 2003; Faria et al., 2011) but many exhibit some hypo-osmoregulatory capacity. In contrast to crab gills (Taylor and Taylor, 1992; Péqueux, 1995; Henry et al., 2012), palaemonid phyllobranchiae do not differ structurally and functionally (Freire et al., 2008a), and participate equally in ion uptake and secretion, ammonia excretion, gas exchange and pH regulation (McNamara et al., 2015). The gill epithelia consist of pillar and septal cells. The apical pillar cell membrane is extensively evaginated as are the intralamellar septal cells, which are rich in mitochondria and connect the bases of adjacent pillar cells (McNamara and Lima, 1997; Belli et al., 2009; McNamara et al., 2015). Salt uptake is the result of transport activity in both ionocyte types. The apical pillar cell membrane holds the V(H^+^)-ATPase (Faleiros et al., 2010), Na^+^ channels and the Cl^−^/HCO_3_^−^ antiporter (McNamara and Faria, 2012) while the Na^+^/K^+^-ATPase predominates in the septal cells (McNamara and Torres, 1999).

Aiming to characterize further the mechanisms that enable salt absorption and secretion by the gills of hololimnetic freshwater crustaceans, we investigate physiological and molecular regulation in *D. pagei* and *M. jelskii* when confronted by osmotic challenge, employing exposure to salt titers lower or higher than those of their natural habitats. Our foremost questions concern whether the osmoregulatory mechanisms of these two phylogenetically distant species that occupy the same osmotic niche depend on the same gill ion transporters, and whether they respond similarly to osmotic challenge.

## Materials and Methods

### Crabs and shrimps

Adult specimens of the semi-terrestrial, freshwater, trichodactylid crab *Dilocarcinus pagei* of either sex, ranging from 3.0 to 6.0 cm in carapace width, were collected from the marginal vegetation of two freshwater reservoirs in Sertãozinho (48° 03’ 06.54” W; 21° 06’ 35.21” S and 48° 03’ 12.57” W; 21° 08’ 26.35” S) in northeastern São Paulo state, Brazil. Specimens of the hololimnetic freshwater shrimp *Macrobrachium jelskii* of either sex, ranging from 3.2 to 5.3 cm total length, were collected from the marginal vegetation of a watercourse at the Araraquara Nautical Club (48° 01’ 33” W; 21° 42’ 17” S), in Americo Brasiliense municipality, central São Paulo state, Brazil. Collections were authorized under SISBIO permit #29594-12 issued by the Brazilian Ministério do Meio Ambiente, Instituto Chico Mendes de Conservação da Biodiversidade (2018-2019).

The animals were kept in large 60-L plastic tanks containing aerated fresh water from their respective collection sites (<0.5 ‰S) for an initial 3-4 day acclimatization period to laboratory conditions (25 °C, natural 14 h light: 10 h dark photoperiod) prior to experimentation. A dry surface in the form of perforated bricks was provided for the crabs while the shrimps were held completely submerged with perforated bricks as a refuge. Diced beef and carrot were offered every two days, and food remnants were removed after a few hours.

### Salinity acclimation

Individual crabs or groups of 6 shrimps each were transferred to aerated 4-L plastic containers respectively filled with 2 or 3 L each of one of two experimental media: either distilled water (0 ‰S), representing a severe hyposmotic challenge, or dilute seawater for the crabs (25 ‰S) and the shrimps (20 ‰S), both salinities constituting a considerable hyperosmotic challenge for each species. The animals were maintained under these conditions for 1, 3, 5, 12 or 24 hours and 3 or 10 days. To avoid the recapture of ions excreted in the urine or lost by diffusion from the body, the distilled water was replaced after 1, 2, 4, 7 and 12 hours and subsequently every 12 hours. The hyperosmotic media were replaced every 24 hours.

The respective control groups consisted of crabs or shrimps maintained under laboratory acclimatization conditions for 7 days, including the initial acclimatization period of 4 days, plus a further 3 days. These groups were considered to represent Time = 0 hours for the respective exposure time courses.

### Osmotic and ionic regulation

After each exposure period, the crabs or shrimps were anesthetized on crushed ice to collect the hemolymph and harvest the gills. For the crabs, a 100-µL hemolymph sample was drawn through a 25-7 gauge needle into a 1-mL plastic syringe via the arthrodial membrane at the junction of the last pereipod and the carapace. Each crab was then bissected and killed, and posterior gill pair #7 was dissected. For the shrimps, a 50-µL hemolymph sample was obtained using an automatic pipette and 10-µL pipette tip inserted into the pericardial sinus at the junction between the cephalothorax and the abdomen. The branchiostegites were then removed and all the gills were dissected.

Hemolymph osmolality (mOsm kg^−1^ H_2_O) in each sample was measured in 10- µL aliquots using a vapor pressure micro-osmometer (Wescor, model 5500, Logan UT). Chloride concentration (mmol L^−1^) was measured in 10-µL aliquots by microtitration against mercury nitrate using s-diphenylcarbazone as an indicator, employing a microtitrator (Metrohm model E 485, Herisau, Switzerland) according to Schales and Schales (1941) adapted by Santos and McNamara (1996). Hemolymph Na^+^ concentration (mmol L^−1^) was measured by atomic absorption spectroscopy (GBC, model 932AA, GBC Scientific Equipment Ltd, Braeside, VIC, Australia) in 5- µL aliquots diluted 1: 20,000 (crabs) or 1: 10,000 (shrimps). Occasionally for the shrimps, a pool of hemolymph from several individuals was used to obtain the volume required.

### RNA extraction and amplification of the gill ribosomal protein L10 and ion transporter partial cDNA sequences

Total RNA was extracted under RNAse-free conditions in Trizol (Invitrogen Corporation, Carlsbad, CA, USA) from posterior gill pair #7 of each crab and from the pooled gills of each shrimp. After DNAse I RNAse-free treatment (Invitrogen) of 1 μg total RNA, mRNA reverse transcription was performed using oligo(dT) primers and Superscript II reverse transcriptase (Invitrogen), according to the manufacturer’s instructions.

Since part of the coding regions for the gill Na^+^/K^+^-ATPase α-subunit (AF409119) and the V(H^+^)-ATPase B-subunit (AF409118) from *D. pagei* were already available from NCBI Genbank (Weihrauch et al., 2004), we designed primers to amplify the partial cDNA sequences for the Na^+^/K^+^/2Cl^−^ symporter (NKCC) and ribosomal protein L10 (RPL10) genes in *D. pagei* (Table 1). For *M. jelskii*, the primers employed successfully to partially amplify the Na^+^/K^+^-ATPase α-subunit, the V(H^+^)-ATPase B-subunit, the NKCC symporter and the RPL10 genes are given in Table 2. All PCR products were verified by electrophoresis in 1% agarose gels.

**Table 1.**
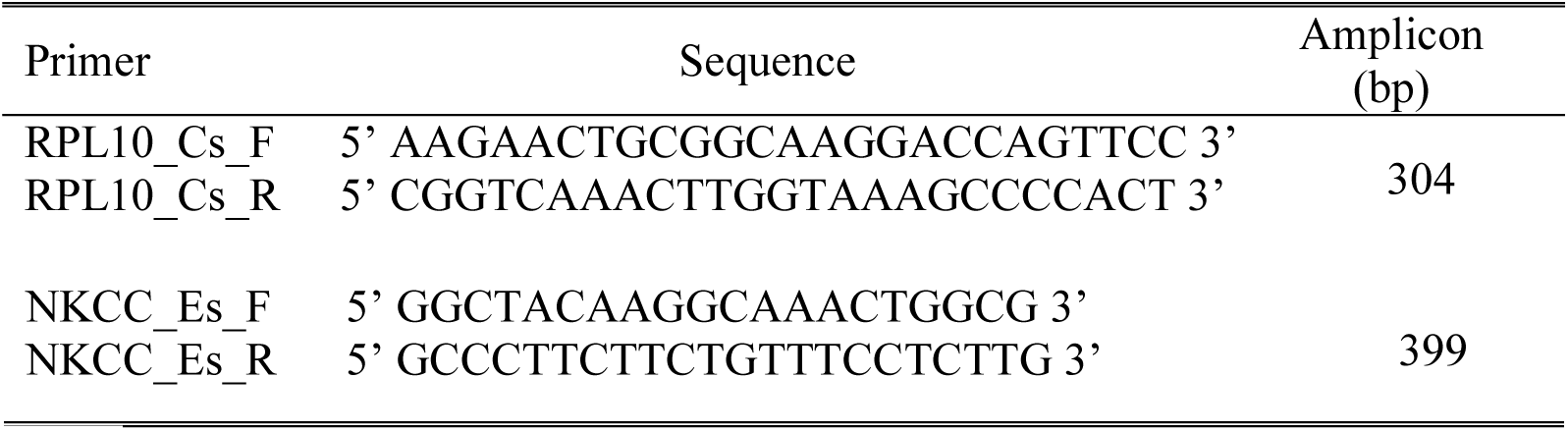
Primer sequences used to amplify partial coding regions of the ribosomal protein L10 gene (RPL10_Cs_F and RPL10_Cs_R) (Wynn et al., 2004) and the Na^+^/K^+^/2Cl^−^ symporter gene (NKCC_Es_F and NKCC_Es_R) in the posterior gills of *Dilocarcinus pagei*.

**Table 2.**
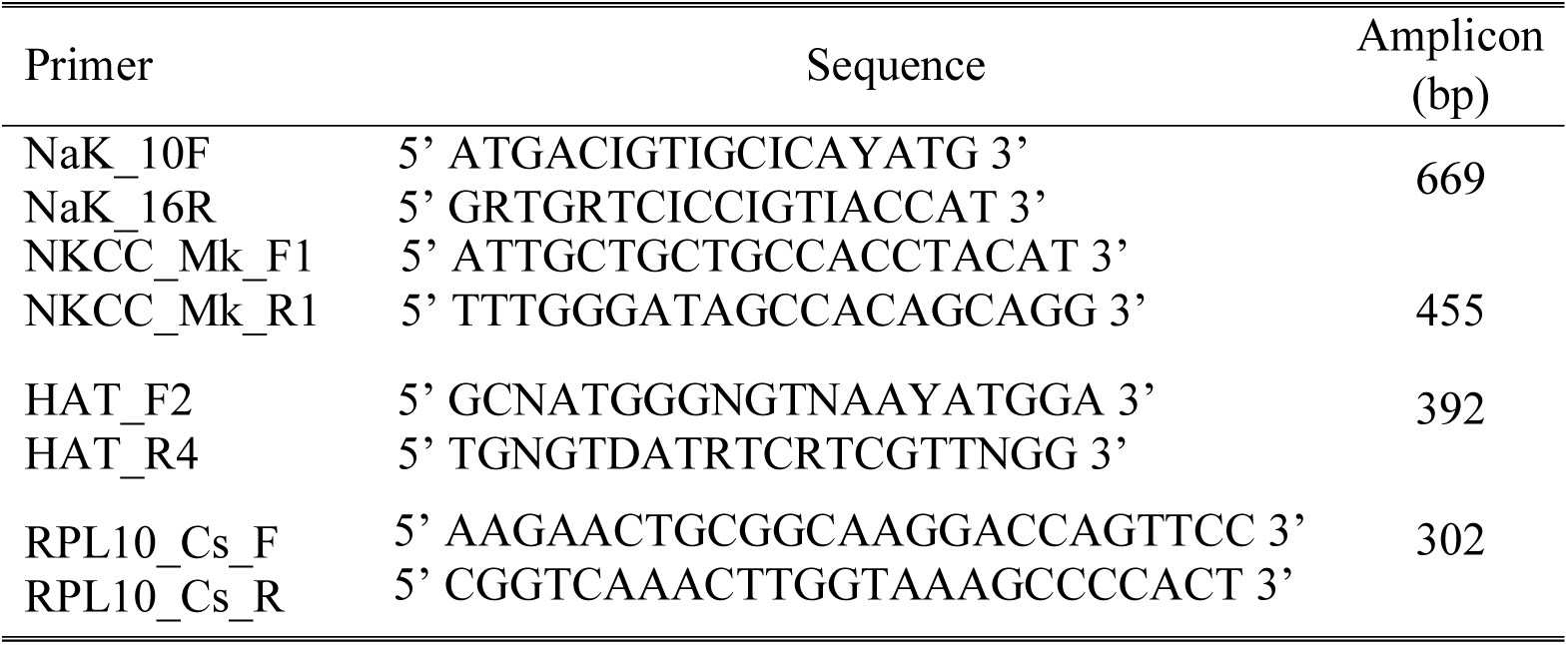
Primer sequences used to amplify partial coding regions of the Na^+^/K^+^- ATPase α-subunit (NaK_10F and NaK_16R) (Towle et al., 2001; Weihrauch et al., 2004), V(H^+^)-ATPase B-subunit (HAT_F2 and HAT_R4) (Weihrauch et al., 2001; Weihrauch et al., 2004), Na^+^/K^+^/2Cl^−^ symporter (NKCC_Mk_F1 and NKCC_Mk_R1) (Rahi et al., 2017) and RPL10 genes (RPL10_Cs_F and RPL10_Cs_R) (Wynn et al., 2004) in the gills of *Macrobrachium jelskii*.

### Cloning and sequencing of the partial cDNA sequences

Individual PCR bands were cut from the agarose gels and the DNA was purified (Qiagen, Valencia, CA, USA). The purified PCR products were cloned into plasmid vectors (pCR 2.1 TOPO TA, Invitrogen, and pJet 1.2/blunt, Thermo Fisher Scientific, USA) that were then used to transform electro-competent DH5α or DH10B Escherichia coli. The recombinant plasmids were extracted with PureLink Quick Plasmid Miniprep Kit (Invitrogen) from successful clones.

The plasmids were sequenced (Genetic Analyzer, ABI PRISM Model 3100, Applied Biosystems, Foster City, CA, USA) using the dideoxynucleotide method (Sanger et al., 1977) employing the forward and reverse primers provided with the cloning kits. Fragment sequences were analyzed for open reading frames (ORF). Searches of GenBank using the BLAST algorithm (Altschul et al., 1990) revealed high similarity with sequences previously published for the coding regions of the genes analyzed in other crustacean species.

The partial cDNA sequences obtained for the *Dilocarcinus pagei* gill RPL10 (GenBank accession number KT876051) and Na^+^/K^+^/2Cl^−^(KX894795) genes, and for the *Macrobrachium jelskii* gill Na^+^/K^+^-ATPase α-subunit (MF615389), V(H^+^)-ATPase B-subunit (MG602347), Na^+^/K^+^/2Cl^−^ symporter (MG566061) and RPL10 (MG566062) genes were then employed to design primers for real-time quantitative gene expression.

Quantitative RT-PCR The relative abundance of target gene mRNA in the total RNA extracts was estimated by quantitative reverse transcription (RT) real-time PCR (StepOnePlus, Applied Biosystems, Carlsbad, CA, USA). Real-time PCR reactions were performed using the Power SYBR Green PCR Master Mix Kit (Applied Biosystems) according to the manufacturer’s instructions employing the primer pairs described in Tables 3 and 4.

**Table 3.**
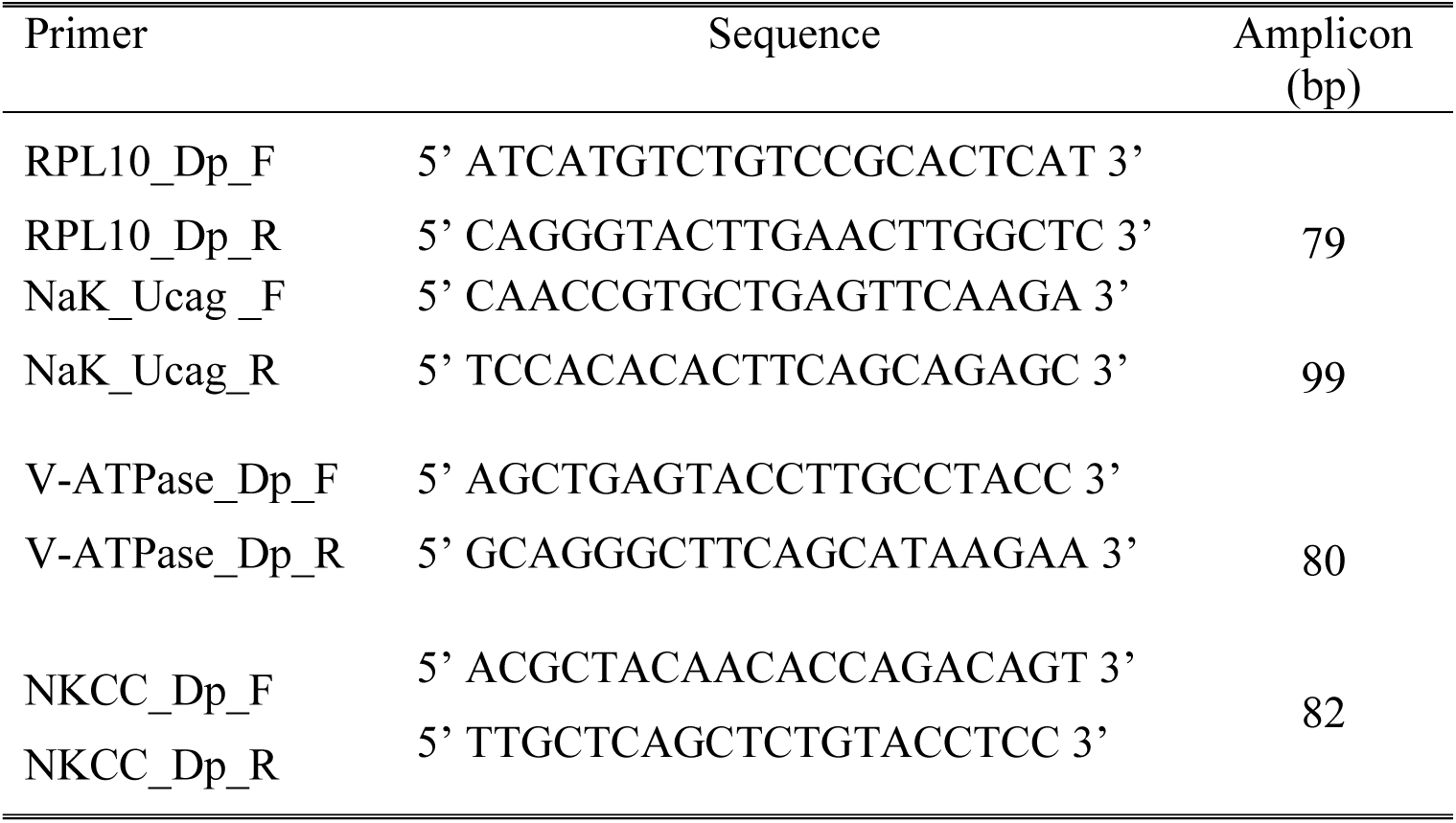
Primer sequences used to quantify mRNA expression of the RPL10 (RPL10_Dp_F and RPL10_Dp_R), Na^+^/K^+^-ATPase α-subunit (NaK_Ucag_F and NaK_Ucag_R), V(H^+^)-ATPase B-subunit (V-ATPase_Dp_F and V-ATPase_Dp_R) and Na^+^/K^+^/2Cl^−^ symporter (NKCC_Dp_F and NKCC_Dp_R) genes in the gills of *Dilocarcinus pagei*.

**Table 4.**
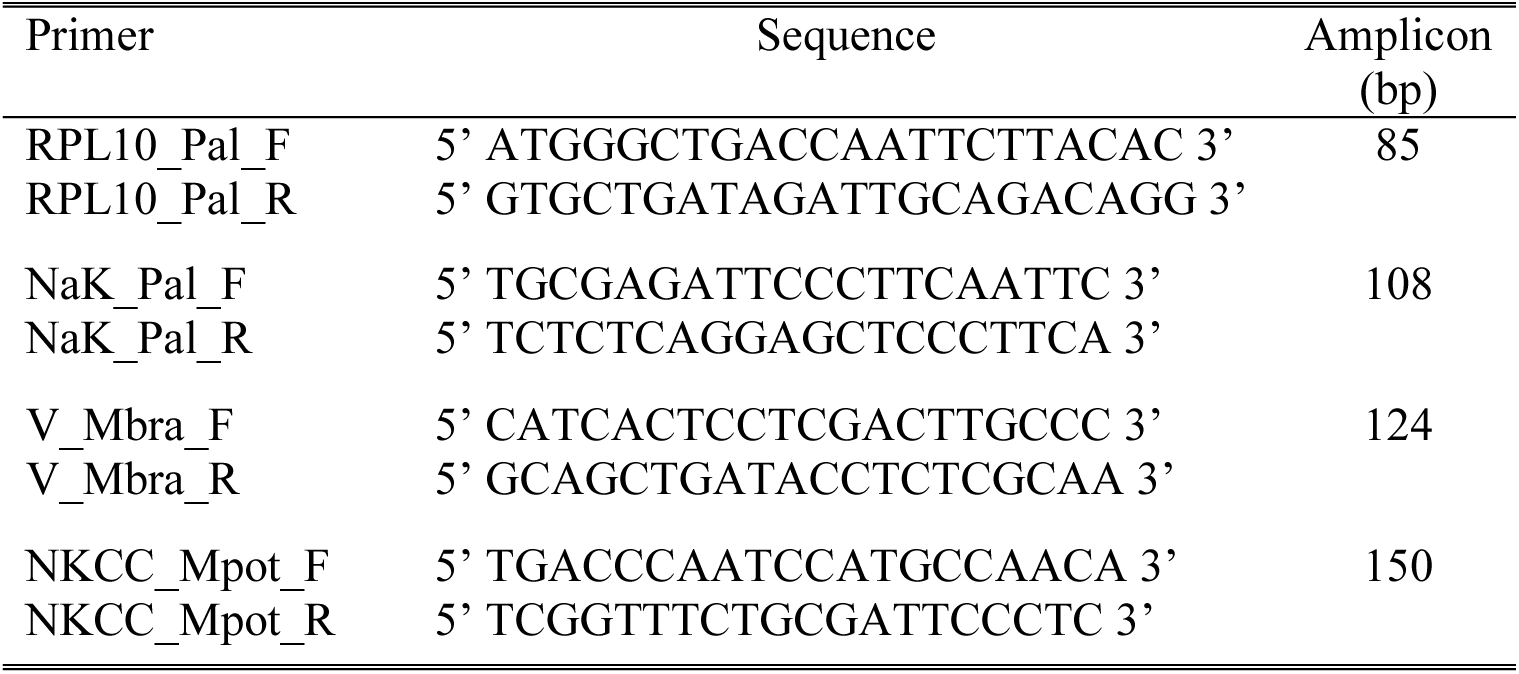
Primer sequences used to quantify mRNA expression of the RPL10 (RPL10_Pal_F and RPL10_Pal_R), Na^+^/K^+^-ATPase α-subunit (NaK_Pal_F and NaK_Pal_R), V(H^+^)-ATPase B-subunit (V_Mbra_F and V_Mbra_R) and Na^+^/K^+^/2Cl^−^ symporter (NKCC_Mpot_F and NKCC_Mpot_R) genes in the gills of *Macrobrachium jelskii*.

The thermocycling procedure consisted of an initial step at 95 °C for 10 min, followed by 40 cycles of 15 s each at 95 °C and a final step at 60 °C for 1 min. The RPL10 gene that encodes for ribosomal protein L10 was used as an endogenous control. Similarity between the amplification efficiencies [E = 10^(−1/slope)^] of the target ion transporter genes and the endogenous RPL10 control gene for each species was evaluated by performing a standard curve validation. Our adjusted curve (R^2^ > 0.99) corresponded to efficiencies between 90 and 110%.

The mRNA expressions of the Na^+^/K^+^-ATPase α-subunit, V(H^+^)-ATPase B-subunit and Na^+^/K^+^/2Cl^−^ symporter were normalized by the expression of the respective ribosomal protein L10 mRNA in the same sample for each species and condition. To compare target gene expression during the time course in each condition (0, and 25 or 20 ‰S) for each species, the normalized data were calibrated by the mean gene expression for the control group (<0.5 ‰S, Time = 0) whose relative arbitrary expression was considered to be ‘1’. To compare gene expressions between *D. pagei* and *M. jelskii* at Time = 0 h (<0.5 ‰S), Time = 12 h (0 ‰S) and Time = 240 h (25 or 20 ‰S), the normalized data were calibrated by the mean expressions of each gene, for each condition, in *D. pagei*. The calibrated data for both comparisons were treated using the exponential formula 2^−∆∆CT^ (Livak e Schmittgen, 2001) and are given as the Mean ± SEM.

### Statistical analyses

After verifying normality of distribution and equality of variance, the data were analyzed using two-way analyses of variance (Species and Exposure Time) to evaluate the main and interactive effects on hemolymph osmolality, chloride and sodium concentrations, and on ion transporter gene expression. Since *M. jelskii* did not survive 24 h exposure to distilled water, the two-way ANOVA was performed only for exposure periods of up to 12 h. For *D. pagei*, a one-way ANOVA (Exposure Time alone) also was performed on data from all exposure periods (0 to 240 h).

When necessary, raw data were normal transformed. However, much of the data concerning transporter gene expression could not be normalized, due to the very marked differences in expression between the two species. In these cases, and for hemolymph [Na^+^] in hyperosmotic media, one-way ANOVAs (Exposure Time alone) were performed since the data were parametric. For exploratory purposes, however, we used two-way ANOVAs, which we considered to be sufficiently robust to provide meaningful effects.

The Student-Newman-Keuls multiple means post-hoc procedure was performed to locate statistically different means. All analyses were performed using SigmaStat 2.03 (Systat Software Inc., San Jose, CA, USA), employing a minimum significance level of P = 0.05. Data are expressed as the Mean ± Standard Error of Mean and were plotted using SlideWrite Plus 4.0 Advanced Graphics Software, Sunnyvale, CA, USA).

To evaluate a species effect on the transcriptional response to salinity challenge, gene expressions of the Na^+^/K^+^-ATPase α-subunit, V(H^+^)-ATPase B subunit and Na^+^/K^+^/2Cl^−^ symporter in *D. pagei* and *M. jelskii* gills also were compared at Time = 0 h (fresh water, <0.5 ‰S), Time = 12 h (0 ‰S) and Time = 240 h (20 or 25 ‰S) using Student’s t-tests (SigmaStat 2.03), employing a minimum significance level of P = 0.05.

## Results

### Time course of changes in hemolymph osmoregulatory parameters during hypoosmotic challenge (0 ‰S)

#### Osmolality

The two-way ANOVA revealed an effect of Exposure Time (F = 3.0, P = 0.026), Species (F = 17.9, P < 0.001) and their interaction (F = 10.3, P = P < 0.001) on hemolymph osmolality when *D. pagei* and *M. jelskii* were exposed to distilled water for up to 12 h. *Macrobrachium jelskii* did not survive more than 12 h in this medium. The one-way ANOVA revealed an effect of Exposure Time on hemolymph osmolality in *D. pagei* (F = 8.1, P < 0.001).

In the control groups (collection site water, <0.5 ‰S, 7 mOsm kg^−1^ H_2_O, Time = 0), hemolymph osmolality was 313.3 ± 22.0 mOsm kg^−1^ H_2_O (N = 7) in *D. pagei*, and 390.0 ± 10.6 mOsm kg^−1^ H_2_O (N = 6) in *M. jelskii* (Figure 1A). In distilled water, hemolymph osmolality increased in *D. pagei* to 396.5 ± 26.2 mOsm kg^−1^ H_2_O (N = 7) after 1 h exposure. However, by 72 h osmolality decreased to 255.4 ± 23.7 mOsm kg^−1^ H_2_O (N = 7), remaining unchanged at around control values up to 240 h. In *M. jelskii*, hemolymph osmolality decreased to 299.7 ± 19.5 mOsm kg^−1^ H_2_O (N = 7) after 12 h exposure.

#### Chloride

The two-way ANOVA revealed an effect of Exposure Time (F = 3.9, P = 0.009), Species (F = 28.3, P < 0.001) and their interaction (F = 8.7, P < 0.001) on hemolymph [Cl^−^] when *D. pagei* and *M. jelskii* were exposed to distilled water for up to 12 h. The one-way ANOVA revealed an effect of Exposure Time on hemolymph [Cl^−^] in *D. pagei* (F = 5.2, P < 0.001).

In the control groups (<0.5 ‰S, 7,7 mmol L^−1^, Time = 0) hemolymph [Cl^−^] was 157.0 ± 13.4 mmol L^−1^ (N = 7) in *D. pagei* and 173.0 ± 6.1 mmol L^−1^ (N = 5) in *M. jelskii* (Figure 1B). In *D. pagei*, hemolymph [Cl^−^] increased 3 (204.1 ± 6.0 mmol L^−1^, N = 5) to 5 h (198.5 ± 9.2 mmol L^−1^, N = 5) after exposure in distilled water, declining to control values by 12 h. Hemolymph [Cl^−^] decreased to 126.3 ± 8.7 mmol L^−1^ (N = 6) in *M. jelskii* exposed for 5 h, and to 107.1 ± 8.9 mmol L^−1^ (N = 3) by 12 h.

#### Sodium

The two-way ANOVA revealed an effect of Exposure Time (F = 9.9, P < 0.001) and Species (F = 55.3, P < 0.001) on hemolymph [Na^+^] in *D. pagei* and *M. jelskii* when exposed for 12 h to distilled water. The one-way ANOVA showed an effect of Exposure Time on hemolymph [Na^+^] in *D. pagei* (F = 6.4, P < 0.001).

In the control groups (<0.5 ‰S, 6.8 mmol L^−1^, Time = 0 h), hemolymph [Na^+^] was 143.0 ± 19.6 mmol L^−1^ (N = 7) in *D. pagei*, and 173.0 ± 10.0 mmol L^−1^ (N = 8) in *M. jelskii* (Figure 1C). In *D. pagei* in distilled water, hemolymph [Na^+^] decreased to 76.5 ± 10.7 mmol L^−1^ (N = 7) by 1 h, remaining unchanged up to 240 h. Hemolymph [Na^+^] in *M. jelskii* decreased in distilled water after 5 h exposure to 123.0 ± 5.4 mmol L^−1^ (N = 7).

**Figure 1.**
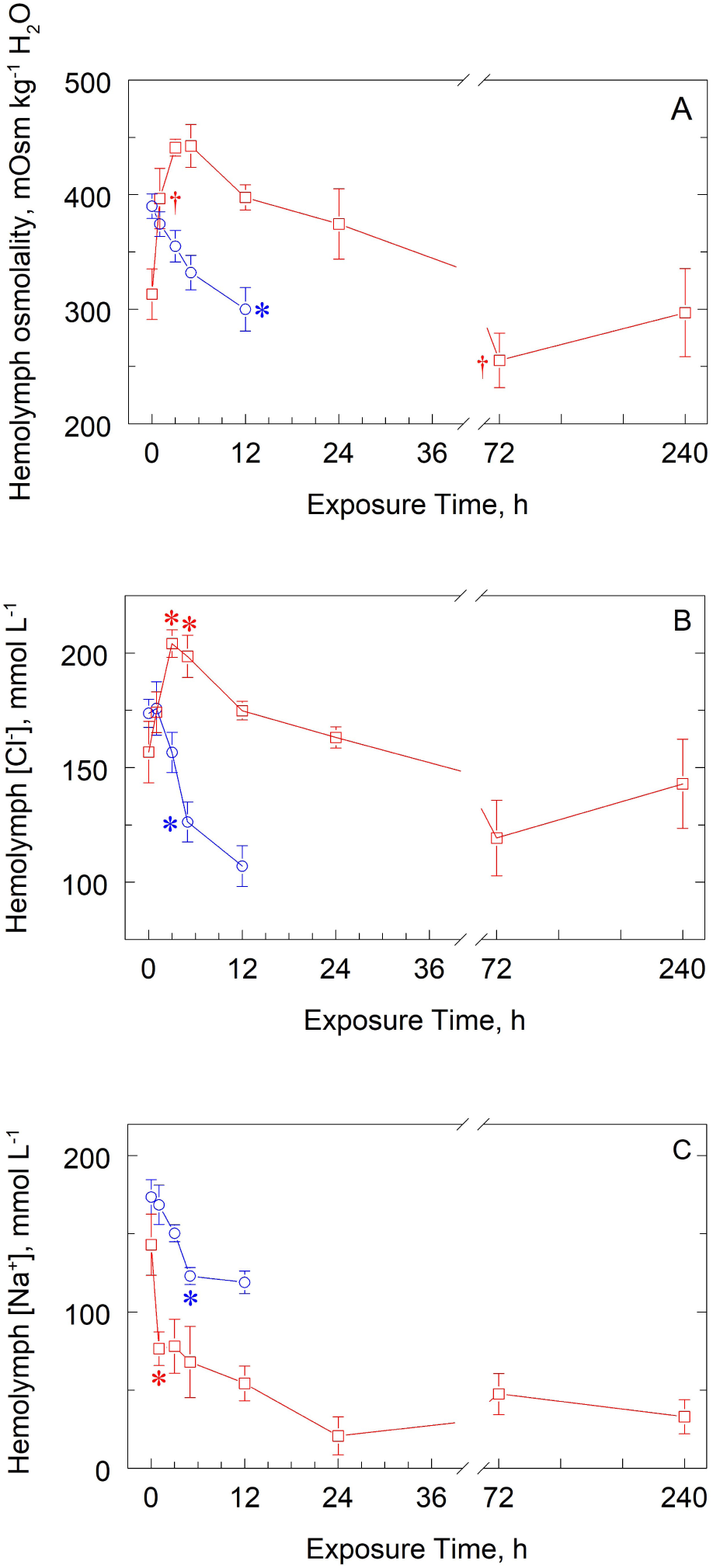
Time courses of changes in osmoregulatory parameters in two hololimnetic decapod crustaceans exposed to a severely hypo-osmotic challenge (distilled water, 0 ‰S) for up to 10 days. A, Hemolymph osmolality (mOsm kg^−1^ H_2_O) in the freshwater crab *Dilocarcinus pagei* (red) and freshwater shrimp *Macrobrachium jelskii* (blue) (Mean ± SEM, 3 ≤ N ≤ 7). B, Hemolymph [Cl^−^] (mmol L^−1^) (Mean ± SEM, 3 ≤ N ≤ 7). C, Hemolymph [Na^+^] (mmol L^−1^) (Mean ± SEM, 5 ≤ N ≤ 8). *Significantly different from the control group (<0.5 ‰S, Time = 0 h). †Significantly different from immediately previous time interval.

### Time course of changes in hemolymph osmoregulatory parameters during hyperosmotic challenge (25 or 20 ‰S)

#### Osmolality

The two-way ANOVA revealed an effect of Exposure Time (F = 49.8, P < 0.001), Species (F = 14.0, P < 0.001) and their interaction (F = 9.6, P < 0.001) on hemolymph osmolality in *D. pagei* and *M. jelskii* when exposed to hyperosmotic media (Figure 2A).

**Figure 2.**
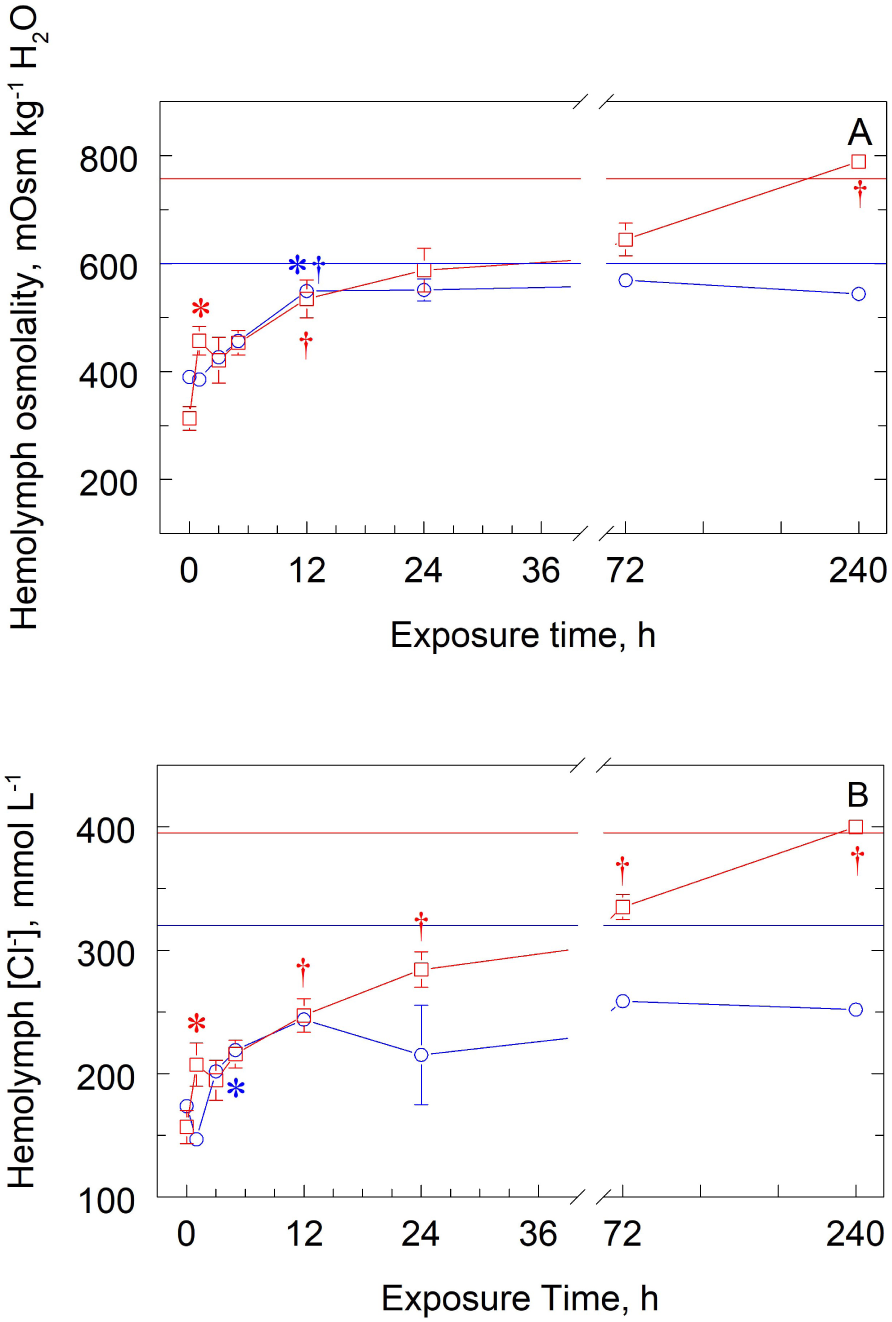
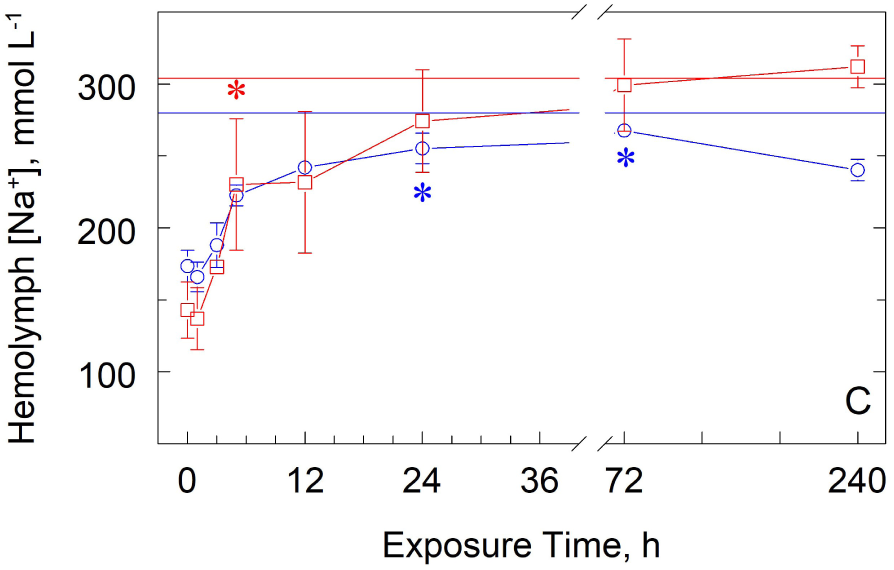
Time courses of changes in osmoregulatory parameters in two hololimnetic decapod crustaceans exposed to a hyper-osmotic challenge (25 or 20 ‰S) for 10 days. A, Hemolymph osmolality (mOsm kg^−1^ H_2_O) in the freshwater crab *Dilocarcinus pagei* (red) and freshwater shrimp *Macrobrachium jelskii* (blue) (Mean ± SEM, 4 ≤ N ≤ 7). B, Hemolymph [Cl^−^] (mmol L^−1^) (Mean ± SEM, 4 ≤ N ≤ 8). C, Hemolymph [Na^+^] (mmol L^−1^) (Mean ± SEM, 5 ≤ N ≤ 9). *Significantly different from the control group (<0.5 ‰S, Time = 0 h). ^†^Significantly different from immediately previous time interval. The dotted red line indicates medium osmolality (757 mOsm kg^−1^ H_2_O), [Cl^−^] (395 mmol L^−1^) and [Na^+^] (304 mmol L^−1^) at 25 ‰S. The dotted blue line indicates medium osmolality (600 mOsm kg^−1^ H_2_O), [Cl^−^] (320 mmol L^−1^) and [Na^+^] (280 mmol L^−1^) at 20 ‰S.

In brackish water (25 ‰S, 757 mOsm kg^−1^ H_2_O), hemolymph osmolality increased in *D. pagei* to 457.0 ± 26.3 mOsm kg^−1^ H_2_O (N = 5) after 1 h and to 534.6 ± 35.1 mOsm kg^−1^ H_2_O (N = 5) by 12 h. After 240 h, osmolality increased further and was approximately isosmotic (789.0 ± 13.3 mOsm kg^−1^ H_2_O, N = 5). In *M. jelskii*, hemolymph osmolality increased to 549.2 ± 12.5 mOsm kg^−1^ H_2_O (N = 6) after 12 h in brackish water (20 ‰S, 600 mOsm kg^−1^ H_2_O) and was slightly hypo-regulated up to 240 h (543.8 ± 3.6 mOsm kg^−1^ H_2_O, N = 5).

#### Chloride

The two-way ANOVA revealed an effect of Exposure Time (F = 45.4, P < 0.001), Species (F = 22.5, P < 0.001) and their interaction (F = 8.0, P < 0.001) on hemolymph [Cl^−^] in *D. pagei* and *M. jelskii* when exposed to hyperosmotic media (Figure 2B).

Hemolymph [Cl^−^] in *D. pagei* in 25 ‰S (395 mmol L^−1^ Cl^−^) increased to 207.2 ± 17.4 mmol L^−1^ (N = 5) after 1 h, and to 335.1 ± 10.1 mmol L^−1^ (N = 7) by 72 h. After 240 h, hemolymph [Cl^−^] was roughly iso-chloremic at 400.0 ± 7.8 mmol L^−1^ (N = 5). In *M. jelskii* in 20 ‰S (320 mmol L^−1^ Cl^−^), hemolymph [Cl^−^] increased to 219.1 ± 6.1 mmol L^−1^ (N = 6) after 5 h exposure, remaining unchanged and clearly hypo- regulated up to 240 h (252.0 ± 7.2 mmol L^−1^, N=6).

#### Sodium

Exposure Time alone affected hemolymph [Na^+^] in *D. pagei* (2-way ANOVA, F = 5.66, P < 0.001) and *M. jelskii* (F = 16.58, P < 0.001) (Figure 2C), both species responding very similarly to hyperosmotic challenge.

In 25 ‰S (304 mmol L^−1^), hemolymph [Na^+^] in *D. pagei* increased after 5 h (230.3 ± 45.7 mmol L^−1^, N = 5) and by 240 h (312 ± 14.5 mmol L^−1^, N = 5) was roughly iso-natriuremic. Hemolymph [Na^+^] in *M. jelskii* increased after 24 h (255.3 ± 10.7 mmol L^−1^, N = 6) and 72 h (267.79 ± 5.9 mmol L^−1^, N = 8) in 20 ‰S, and was clearly hypo-regulated.

### Ionic contribution to hemolymph osmolality after hypo-osmotic (0 ‰S) or hyperosmotic challenge (20 or 25 ‰S)

Control *D. pagei* (<0.5 ‰S, Time = 0 h) showed a [Cl^−^]: osmolality ratio of 0.50: 1 and a [Na^+^]: osmolality ratio of 0.46: 1. In control *M. jelskii* (<0.5 ‰S, Time = 0 h), both ratios were 0.44: 1.

After 10-days acclimation of *D. pagei* in 0 ‰S, these ratios were 0.48: 1 for [Cl^−^] and 0.10: 1 for [Na^+^]. In *M. jelskii* exposed for 12 h, ratios were 0.36: 1 for [Cl^−^] and 0.40: 1 for [Na^+^].

In hyperosmotic media after 10 days, these ratios were 0.51: 1 for [Cl^−^] and 0.40: 1 for [Na^+^] in *D. pagei*, and 0.46: 1 and 0.44: 1 in *M. jelskii*.

### Time course of changes in ion transporter gene expression during hyposmotic challenge (0 ‰S)

#### V(H^+^)-ATPase

The two-way ANOVA revealed an effect of Exposure Time (F = 3.3, P = 0.019) and Species (F = 18.8, P < 0.001) but not their interaction (F = 1.9, P = 0.127) on V(H^+^)-ATPase B-subunit expression in the gills of *D. pagei* and *M. jelskii* when exposed to distilled water for up to 12 h. The one-way ANOVA revealed an absence of effect of Exposure Time on gill V(H^+^)-ATPase gene expression in *D. pagei* (F = 1.8, P = 0.112) (Figure 3A).

**Figure 3.**
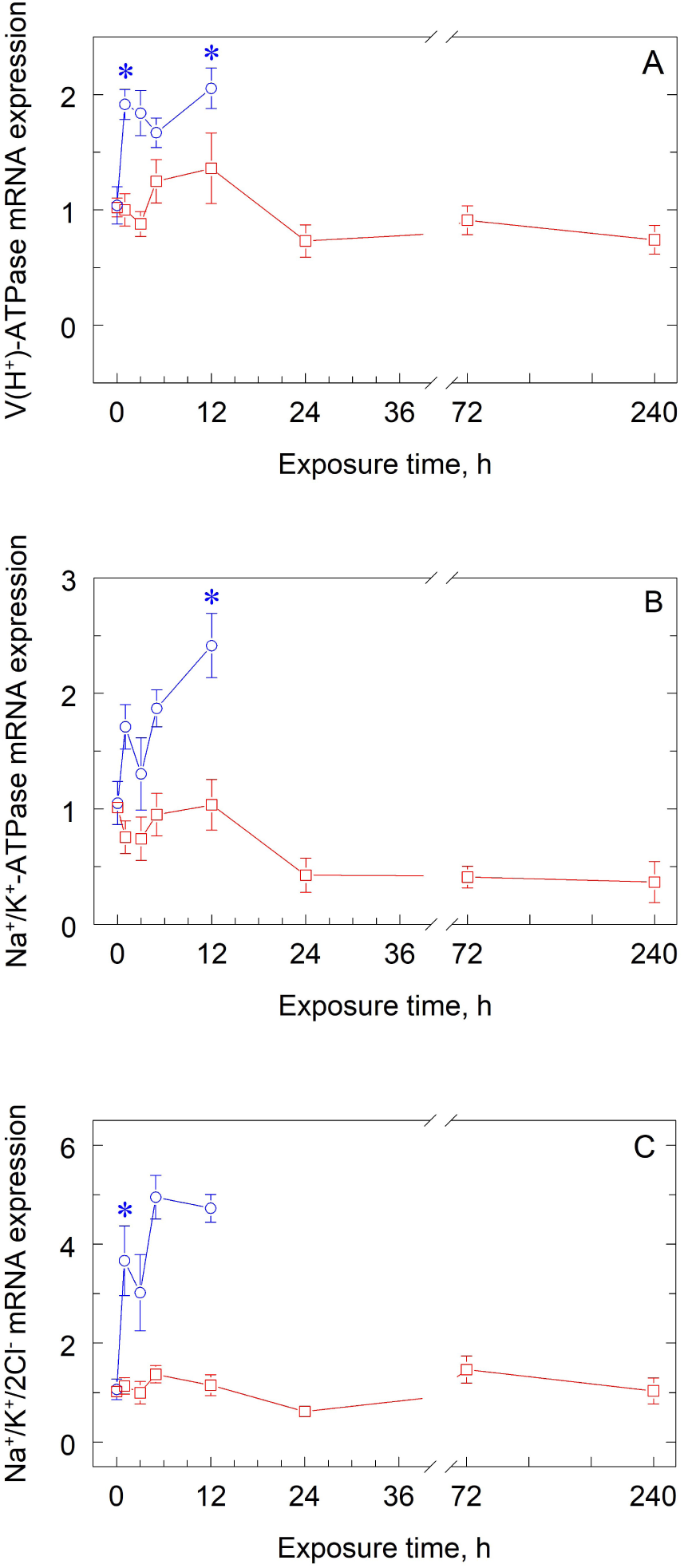
Time courses of changes in ion transporter gene expression in the gills of two hololimnetic decapod crustaceans exposed to a severely hypo-osmotic challenge (distilled water, 0 ‰oS) for up to 10 days. A, V(H^+^)-ATPase B-subunit gene expression in the freshwater crab *Dilocarcinus pagei* (red) and freshwater shrimp *Macrobrachium jelskii* (blue) (Mean ± SEM, 4 ≤ N ≤ 7). B, Na^+^/K^+^-ATPase α-subunit gene expression (Mean ± SEM, 4 ≤ N ≤ 7). C, Na^+^/K^+^/2Cl^−^ symporter gene expression (Mean ± SEM, 4 ≤ N ≤ 7). *Significantly different from the control group (<0.5 ‰S, Time = 0 h). Target gene mRNA expression has been normalized against expression of the respective ribosomal protein L10 in the same sample and calibrated against expression in the control group at Time = 0 h.

There were no changes in gill V(H^+^)-ATPase B-subunit expression in *D. pagei* during the 10-day time course (Figure 3A). In *M. jelskii*, gill V(H^+^)-ATPase expression increased by 1.9- to 2-fold after 1 and 12 h compared to the control group (<0.5 ‰S, Time = 0, N = 4).

#### Na^+^/K^+^-ATPase

The two-way ANOVA revealed an effect of Exposure Time (F = 4.0, P = 0.007), Species (F = 33.8, P < 0.001) and their interaction (F = 2.9, P = 0.032) on gill Na^+^/K^+^-ATPase α-subunit gene expression in *D. pagei* and *M. jelskii* when exposed to distilled water for up to 12 h. The one-way ANOVA revealed an effect of Exposure Time on gill Na^+^/K^+^-ATPase gene expression in *D. pagei* (F = 3.5, P = 0.005) (Figure 3B).

Gill Na^+^/K^+^-ATPase α-subunit gene expression was unchanged over the entire 10-day time course of exposure to distilled water in *D. pagei* (Figure 3B). In *M. jelskii*, gill Na^+^/K^+^-ATPase expression increased 2.4-fold after 12 h compared to control group (<0.5 ‰S, Time = 0, N = 4).

#### Na^+^/K^+^/2Cl^−^ symporter

The two-way ANOVA revealed an effect of Exposure Time (F = 7.4, P < 0.001), Species (F = 76.9, P < 0.001) and their interaction (F = 5.6, P = 0.032) on gill Na^+^/K^+^/2Cl^−^ symporter gene expression in *D. pagei* and *M. jelskii* exposed to distilled water for up to 12 h. The one-way ANOVA revealed an absence of effect of Exposure Time on Na^+^/K^+^/2Cl^−^ gene expression in *D. pagei* (F = 1.5, P = 0.186) (Figure 3C).

There were no changes in gill Na^+^;K^+^/2Cl^−^ gene expression in *D. pagei*. In *M. jelskii*, gill Na^+^/K^+^/2Cl^−^ gene expression increased by 3- to 5-fold from 1 h exposure onwards (Figure 3C).

### Time course of changes in ion transporter gene expression during hyperosmotic challenge (25 or 20 ‰S)

#### V(H^+^)-ATPase

The two-way ANOVA revealed an effect of Exposure Time (F = 7.8, P < 0.001), Species (F = 217.8, P < 0.001) and their interaction (F = 8.2, P < 0.001) on V(H^+^)-ATPase B-subunit gene expression in the gills of *D. pagei* and *M. jelskii* when exposed to hyperosmotic media (Figure 4A).

In 25 ‰S, gill V(H^+^)-ATPase B-subunit gene expression decreased in D. pagei to from 0.23- to 0.15-fold after 72 to 240 h exposure compared to the control group (<0.5 ‰S, Time = 0, N = 4). In *M. jelskii*, V(H^+^)-ATPase gene expression increased 2- to 3-fold after 3 to 12 h exposure at 20 ‰S, declining to control values by 24 h (Figure 4A).

**Figure 4.**
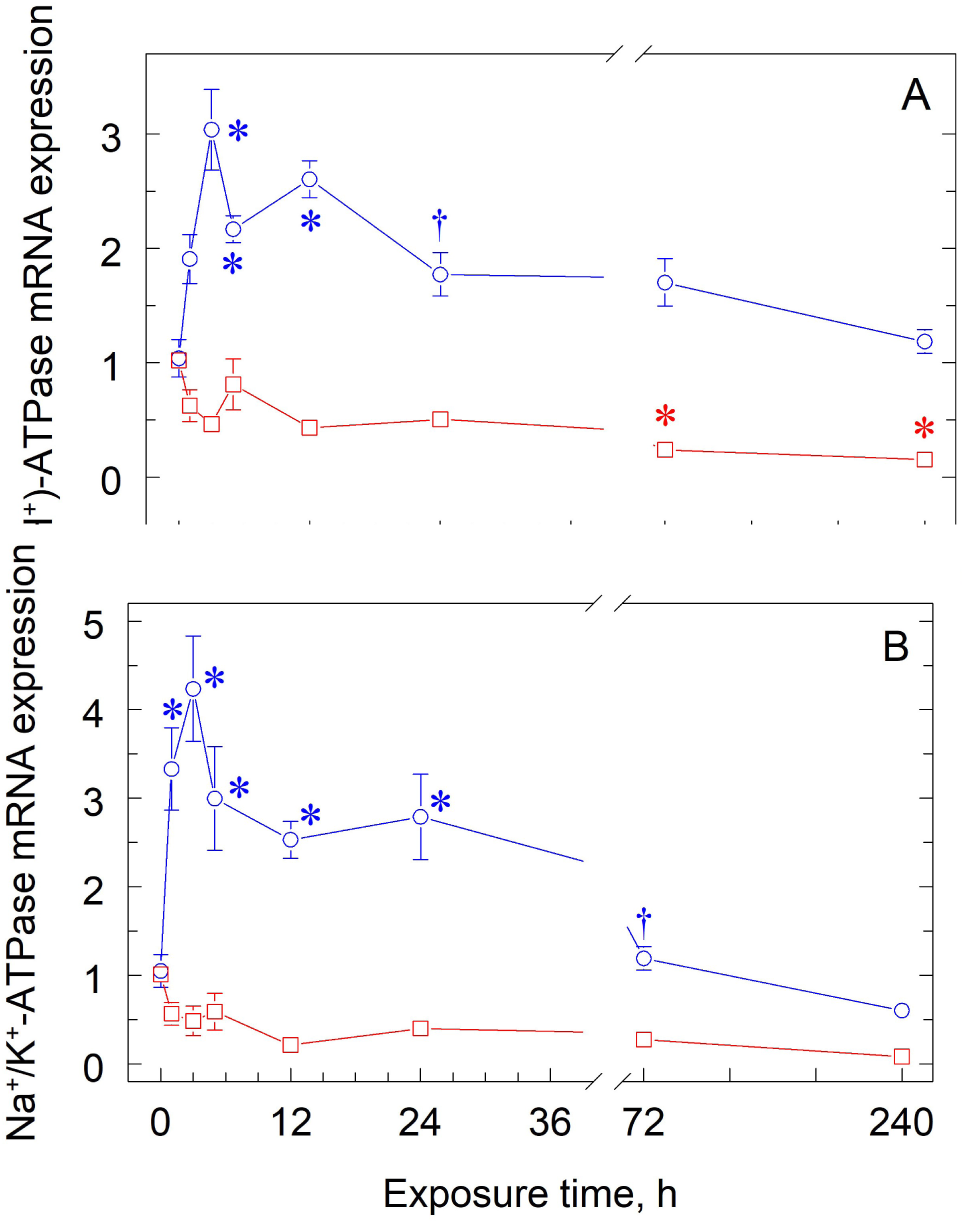
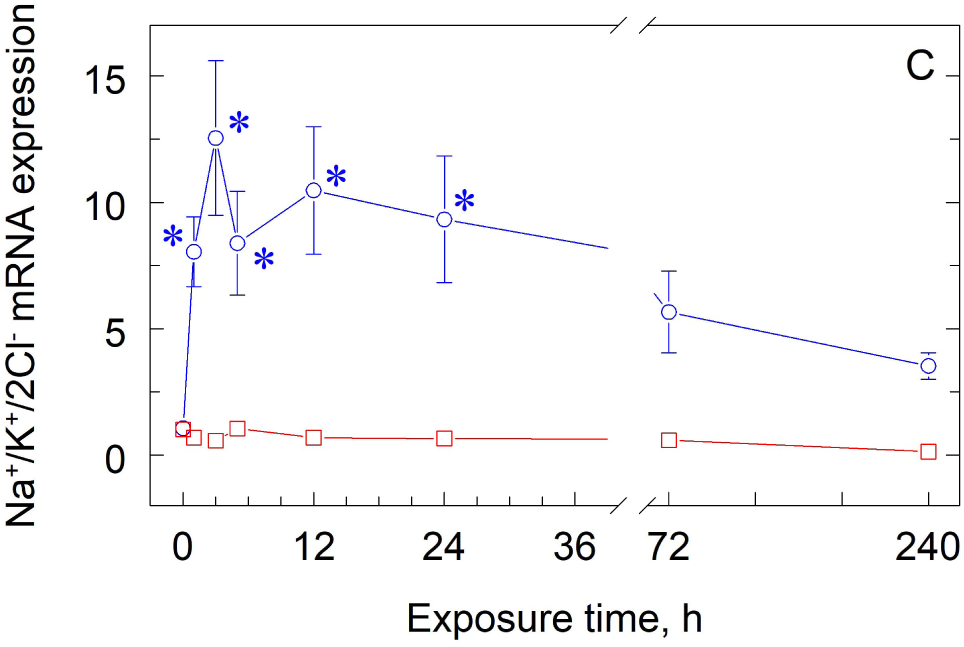
Time courses of changes in ion transporter gene expression in the gills of two hololimnetic decapod crustaceans exposed to a hyperosmotic challenge (25 or 20 ‰S) for 10 days. A, V(H^+^)-ATPase B-subunit gene expression in the freshwater crab *Dilocarcinus pagei* (red) and freshwater shrimp *Macrobrachium jelskii* (blue) (Mean ± SEM, 4 ≤ N ≤ 7). B, Na^+^/K^+^-ATPase α-subunit gene expression (Mean ± SEM, 4 ≤ N ≤ 7). C, Na^+^/K^+^/2Cl^−^ symporter gene expression (Mean ± SEM, 4 ≤ N ≤ 7). *Significantly different from the control group (<0.5 ‰S, Time = 0 h). ^†^Significantly different from immediately previous time interval. Target gene mRNA expression has been normalized against expression of the respective ribosomal protein L10 in the same sample and calibrated against expression in the control group at Time = 0 h.

#### Na^+^/K^+^-ATPase

The two-way ANOVA revealed an effect of Exposure Time (F = 9.2, P < 0.001), Species (F = 137.0, P < 0.001) and their interaction (F = 7.9, P < 0.001) on Na^+^/K^+^-ATPase α-subunit gene expression in the gills of *D. pagei* and *M. jelskii* when exposed to hyperosmotic media (Figure 4B).

Gill Na^+^/K^+^-ATPase gene expression was unchanged during the 10-day time course of exposure to 25 ‰S in *D. pagei*. In *M. jelskii*, gill Na^+^/K^+^-ATPase gene expression increased ≈3- to 4-fold between 1 to 24 h exposure at 20 ‰S. After 72 h exposure, expression returned to control levels (Figure 4B).

#### Na^+^/K^+^/2Cl^−^ symporter

The two-way ANOVA revealed an effect of Exposure Time (F = 3.0, P = 0.009), Species (F = 74.2, P < 0.001) and their interaction (F = 2.9, P = 0.009) on Na^+^/K^+^/2Cl^−^ symporter gene expression in the gills of *D. pagei* and *M. jelskii* when exposed to hyperosmotic media (Figure 4C).

Gill Na^+^/K^+^/2Cl^−^ symporter gene expression was unchanged in *D. pagei* after 10 days exposure in 25 ‰S. In *M. jelskii*, gill Na^+^/K^+^/2Cl^−^ gene expression increased 3- to 12.5-fold between 1 and 24 h exposure, declining to control levels by 72 h and thereafter (Figure 4C).

### Interspecific comparison of gill ion transporter gene expression

Quantitative data for the gene expressions of the V(H^+^)-ATPase B-subunit, Na^+^/K^+^-ATPase α-subunit and Na^+^/K^+^/2Cl^−^ symporter in the gills of *D. pagei* and *M. jelskii* at relevant points during the acclimation time course are provided in Table 5. Calibrating the expressions for *M. jelskii* against those for *D. pagei* allows comparison of the unacclimated, basal transcription rates and of the responses to osmoregulatory challenge in the two species.

**Table 5.** Relative quantitation of gene expressions of the Na^+^/K^+^-ATPase α-subunit, V(H^+^)-ATPase B subunit and Na^+^/K^+^/2Cl^−^ symporter in the gills of *Dilocarcinus pagei* and *Macrobrachium jelskii*, at selected intervals during the time course of osmoregulatory acclimation. Data are given for unacclimated control animals (Time = 0 h, <0.5 ‰S, fresh water), for severe hypo-osmoregulatory challenge at Time = 12 h (0 ‰S, distilled water) and for hyper-osmoregulatory challenge at Time = 240 h (25 or 20 ‰S, respectively). Gene expressions for *M. jelskii* are further calibrated against the respective expressions for *D. pagei* at each time interval and condition analyzed. Data are the Mean ± SEM (4 ≤ N ≤ 7). *Significantly different from respective gene and condition in *D. pagei*.

|  |  | V(H<sup>+</sup>)-ATPase | Na<sup>+</sup>/K<sup>+</sup>-ATPase | Na<sup>+</sup>/K<sup>+</sup>/2Cl<sup>-</sup> |
| --- | --- | --- | --- | --- |
| *D. pagei* | Control, <0.5 ‰S | 1.02 ± 0.08 | 1.01 ± 0.07 | 1.02 ± 0.09 |
|  | 12 h, 0 ‰S | 1.11 ± 0.25 | 1.08 ± 0.23 | 1.07 ± 0.20 |
|  | 240 h, 25 ‰S | 1.09 ± 0.26 | 1.41 ± 0.66 | 1.50 ± 0.79 |
| *M. jelskii* | Control, <0.5 ‰S | *0.51 ± 0.08 | *4.48 ± 0.80 | 0.81 ± 0.16 |
|  | 12 h, 0 ‰S | 0.82 ± 0.07 | *11.89 ± 1.37 | *3.35 ± 0.20 |
|  | 240 h, 20 ‰S | *4.12 ± 0.36 | *49.49 ± 8.22 | *28.47 ± 4.25 |

In the control, acclimatized animals in fresh water (Time = 0 h, <0.5 ‰S), V(H^+^)-ATPase gene expression was ≈0.5-fold less (P = 0.003) and Na^+^/K^+^-ATPase expression was ≈4-fold greater (P < 0.001) in *M. jelskii*. Na^+^/K^+^/2Cl^−^ symporter gene expressions were similar (P = 0.227). After severe hypo-osmotic challenge for 12 h in distilled water (0 ‰S), the Na^+^/K^+^-ATPase and Na^+^/K^+^/2Cl^−^ gene expressions were respectively ≈12- and ≈3-fold greater (P < 0.001) in *M. jelskii*. V(H^+^)-ATPase gene expressions were similar (P = 0.220). After hyper-osmoregulatory challenge for 240 h (25 or 20 ‰S), all gene expressions were considerably greater (P < 0.001) in *M. jelskii*, reaching ≈4-fold for the V(H^+^)-ATPase, ≈49-fold for the Na^+^/K^+^-ATPase and ≈28-fold for the Na^+^/K^+^/2Cl^−^ symporter.

## Discussion

This investigation examined patterns of change in osmoregulatory parameters like hemolymph osmolytes and gill ion-transporter gene expression, during exposure of two phylogenetically unrelated, hololimnetic crustaceans from very similar habitats to extremely dilute, or concentrated media. Virtually all the response parameters differ markedly between the two species. *Dilocarcinus pagei* is an old (≈80 million years) (Collins et al., 2011; Tsang et al., 2014), well-adapted freshwater crab (Augusto et al., 2007b) while species of the genus Macrobrachium, like the freshwater shrimp *M. jelskii*, are more recent inhabitants (≈30 million years) (Short, 2004; Murphy and Austin, 2005; De Grave et al., 2009; Ashelby et al., 2012). This difference in evolutionary time in fresh water may underlie many of their differences in response to osmotic challenge.

### Osmotic and ionic regulation

*Dilocarcinus pagei* and *Macrobrachium jelskii* clearly employ two distinct osmoregulatory strategies: on extended, severe hypo-osmotic challenge, the crab maintains its osmotic and ionic regulatory ability while the shrimp steadily loses hemolymph ions and succumbs to osmotic dilution within 12 h of exposure. On hyper-osmotic challenge, *D. pagei* tolerates elevated extra- and intracellular isosmoticity, while *M. jelskii* effects anisosmotic and anisoionic extracellular regulation.

Fresh caught *D. pagei* show [Cl^−^]: and [Na^+^]: osmolality ratios of 0.50: 1 and 0.46: 1, respectively, similar to previous findings (0.53-0.54: 1 and 0.49-0.53: 1) (Augusto et. al., 2007b; Onken and McNamara, 2002). Slightly lesser ratios (both 0.44: 1) characterize the hemolymph of *M. jelskii*. Thus, [Cl^−^] and [Na^+^] account for ≈97% of hemolymph osmolality, suggesting a very minor contribution of organic osmolytes such as free amino acids (FAA). This highlights the role of powerful mechanisms of anisosmotic extracellular regulation in maintaining the osmolality, ionic concentration and volume of the hemolymph in these two freshwater species in their natural habitat, likely owing to the activity of ion-transporting proteins like the Na^+^/K^+^-ATPase, V(H^+^)-ATPase, the Na^+^/K^+^/2Cl^−^ symporter and ion channels (McNamara and Faria, 2012).

*Dilocarcinus pagei* when challenged with a severely hyposmotic medium, unexpectedly exhibits transiently increased hemolymph osmolality and [Cl^−^] that subsequently decline to control levels. In contrast, [Na^+^] diminishes very rapidly, attaining a stable, low value around ≈50 mmol L^−1^. After 10-days acclimation, [Cl^−^] is well regulated with a [Cl^−^]: osmolality ratio of 0.48: 1, similar to fresh caught crabs. However, [Na^+^] is regulated at ≈30% of control titers, the [Na^+^]: osmolality ratio reaching just 0.10: 1, suggesting the presence of a compensatory osmolyte like NH_4_^+^ that might buffer hemolymph osmolality. Total FAA concentrations contribute very little to hemolymph osmolality in *D. pagei* when in fresh water (0.48 ± 0.06 mmol/L, Augusto et al., 2007b).

In contrast, *M. jelskii* exhibits both 12-h [Cl^−^]: and [Na^+^]: osmolality ratios (0.36: 1 and 0.40, respectively) similar to control shrimps. However, hemolymph osmolality, [Cl^−^] and [Na^+^] also decline by ≈30%, incompatible with survival beyond 12 h in distilled water, despite the species ability to hyper-regulate well in fresh water (44: 1). This lack of tolerance may derive from disproportionate water influx and ion loss owing to its large surface/volume ratio compared to *D. pagei*. Caridean shrimps exhibit a cylindrical body form and a well-developed abdomen. Brachyuran crabs show a compact body morphology owing to their small, ventrally folded abdomen, which reduces the area available for passive water and ion fluxes (Ruppert and Barnes, 1994; Schmidt-Nielsen, 2002).

In *D. pagei* exposed to a sub-lethal hyperosmotic medium, hemolymph osmolality, [Cl^−^] and [Na^+^] increase gradually over the 10-day acclimation period, becoming isosmotic, iso-chloremic and iso-natriuremic, revealing the absence of ion- specific hypo-regulatory mechanisms. Augusto et al. (2007b) have disclosed a more rapid, two-day time course for osmotic and ionic adjustment. The 10-day [Cl^−^]: and [Na^+^]: osmolality ratios are mainly unchanged (0.51: 1 and 0.40: 1, respectively), revealing balanced regulation. Similar ratios calculated from Augusto et al. (2007b) (0.47: 1 and 0.67: 1, respectively), suggest a greater contribution of [Na^+^] to osmolality. Despite a minimal contribution to hemolymph osmolality (≈1 mOsm kg^−1^ H_2_O), FAA synthesis and hemolymph or muscle protein catabolism follow a much slower time course (≈10 days, Augusto et al., 2007b). Clearly, *D. pagei* has maintained the capacity to tolerate a very elevated cellular isosmoticity (≈800 mOsm kg^−1^ H_2_O) despite its lengthy evolutionary history in fresh water, a possible trade-off against the inability to secrete salt and its ensuing adaptive value (McNamara and Faria, 2012; McNamara et al., 2015).

The exposure of *M. jelskii* to a sub-lethal hyperosmotic medium reveals that while hemolymph osmolality, [Cl^−^] and [Na^+^] increase gradually during the 10-day acclimation period, these parameters are clearly hypo-regulated at 80-90% of ambient titers. The [Cl^−^]: and [Na^+^]: to osmolality ratios (0.46: 1 and 0.44: 1, respectively) are unchanged, revealing balanced ionic regulation and a minor contribution of organic osmolytes to hemolymph osmolality. Macrobrachium amazonicum from a land- locked freshwater population show [Cl^−^]: and [Na^+^]: to osmolality ratios of 0.37 and 0.30, respectively, shifting to 0.42 and 0.59 after 10 days at 25 ‰S (Augusto et al., 2007a). Thus, organic osmolytes contribute little to hemolymph osmolality in *M. jelskii* and M. amazonicum, revealing balanced ionic regulation via anisosmotic extracellular mechanisms. Like *M. jelskii*, palaemonids from fluctuating salinity regimes such as M. equidens (Denne, 1968), M. olfersii and Palaemon pandaliformis (Freire et al., 2003) and *M. acanthurus* and P. northropi (Faleiros et al., 2017) exhibit distinct hypo-osmoregulatory ability.

### Gill ion-transporter gene expression

Gill ion-transporter gene expression differs markedly between the two species under both hypo- and hyper-osmotic challenges.

In *D. pagei* in hyposmotic medium, transporter mRNA expression of the main gill ion-transporter genes, i. e., the V(H^+^)-ATPase B-subunit, Na^+^/K^+^-ATPase α- subunit and the Na^+^/K^+^/2Cl^−^ symporter, was unaltered over the entire acclimation period. Nevertheless, hemolymph osmolality and [Cl^−^] were essentially well-regulated in the long run; hemolymph [Na^+^] alone declined drastically. This suggests that the basal rates of transporter gene transcription measured at the outset of hyposmotic challenge are sufficient to sustain the strong hyper-regulatory ability of *D. pagei* seen after 10 days acclimation in distilled water. Clearly, the kinetic characteristics of these transporter enzymes, particularly the V(H^+^)- (K_M_ = 4.2 ± 0.3 mmol L^−1^ in fresh water, Firmino et al., 2011) and Na^+^/K^+^-ATPases (K_0.5_ = 0.34 ± 0.02 μmol L^−1^ at the high- affinity sites and K_M_ = 84.0 ± 5.0 μmol L^−1^ at the low-affinity sites, Furriel et al., 2010), must enable ion uptake sufficient to sustain the enormous gradient imposed.

In *M. jelskii*, however, mRNA expression of the gill V(H^+^)-ATPase B subunit, Na^+^/K^+^-ATPase α-subunit and Na^+^/K^+^/2Cl^−^ symporter increased from 2 to 5-fold on 12-h hyposmotic challenge, showing that gene transcription of these transporters is activated by the ion poor medium. Even so, in stark contrast to *D. pagei*, these augmented transporter transcription rates were insufficient to maintain the osmotic and ionic gradients necessary to sustain systemic integrity, leading to death. Gill V(H^+^)-ATPase B subunit transcription rate likewise increases 10-fold in the tidepool shrimp *Palaemon northropi* faced with a 10-day osmotic challenge in dilute seawater (8 ‰S) (Faleiros et al., 2017). Interestingly, the 5-fold increase in Na^+^/K^+^/2Cl^−^ symporter expression in *M. jelskii* suggests a role for this transporter in salt uptake in fresh water. This symporter has been suggested previously to participate only in salt uptake in hyper-regulating, brackish water crustaceans (Faleiros et al., 2010; Towle et al., 2011; Henry et al., 2012; Havird et al., 2014; Boudour-Boucheker et al., 2016).

In hyperosmotic medium, gill ion-transporter mRNA expression in *D. pagei* either decreases slightly, as seen for the V(H^+^)-ATPase B-subunit, or remains unaltered as in the Na^+^/K^+^-ATPase α-subunit and the Na^+^/K^+^/2Cl^−^ symporter gene. Active participation of these transporters in anisosmotic regulation seems unlikely, since *D. pagei* does not hypo-regulate, becoming isosmotic and iso-ionic. A direct role for the V(H^+^)-ATPase in hypo-regulatory adjustment is thus unlikely, which is corroborated by the rapid time- (1 h in 21 ‰S) and salinity- (5 to 21 ‰S for 10 days) dependent reduction in V(H^+^)-ATPase phosphohydrolytic activity seen in *D. pagei* posterior gills on salinity challenge, Firmino et al., 2011). This is an important transport decoupling component of adjustment to salinity, since active, V(H^+^)- ATPase-dependent Na^+^ uptake will rapidly decrease, and intracellular H^+^ accumulation will affect pH balance, kinetics of CO^2^ hydration by carbonic anhydrase, and apical Cl^−^/HCO_3_^−^ exchange. The unaltered expression of Na^+^/K^+^- ATPase α-subunit mRNA suggests that basal transcription rates are sufficient for housekeeping purposes in the ionocytes. The constant low expression of the gill Na^+^/K^+^/2Cl^−^ symporter gene underscores the crab’s inability to hypo-regulate Cl^−^ and Na^+^, the evolutionary option having been reliance on a strategy of isosmotic intracellular adjustment, likely maintained in part by the symporter (Amado et al., 2006; Foster et al., 2010; Freire et al., 2013). Like *D. pagei*, the hololimnetic freshwater anomuran Aegla schmitti shows decreased (≈50%) gill V(H^+^)-ATPase activity after 10 days acclimation to 25 ‰S while Na^+^/K^+^-ATPase and carbonic anhydrase activities are unchanged (Bozza et al., 2019).

In *M. jelskii* acclimated to hyperosmotic medium, mRNA expression of all three gill ion transporter genes increases markedly although transiently, returning to basal levels by 72 h. While the V(H^+^)-ATPase may not participate in salt secretion in some palaemonids and hermit crabs (Faleiros et al., 2010, 2018) or in *D. pagei* (Firmino et al., 2011), V(H^+^)-ATPase B subunit expression does increase in high salinity media in the estuarine crab *Neohelice granulata* (Luquet et al., 2005). The V(H^+^)-ATPase also drives gill NH_4_^+^ excretion in crabs (Weihrauch et al., 2002, 2004), and increased transcription may reflect this demand. An indirect role for the V(H^+^)-ATPase in salt secretion in the Crustacea warrants wider investigation.

Gill Na^+^/K^+^-ATPase α-subunit expression increases initially in *M. jelskii* as seen in *Palaemon northropi* after 5 days acclimation to concentrated seawater (50 ‰S), but not in *M. acanthurus* (25 ‰S) (Faleiros et al., 2017). Increased Na^+^/K^+^- ATPase expression likely subsidizes increased Na^+^ and K^+^ turnover through the basal Na^+^/K^+^/2Cl^−^ symporter (McNamara and Faria, 2012) and may thus regulate Cl^−^ transport. The coupled activity of these two transporters should be examined more closely. To illustrate, mRNA expression of the Na^+^/K^+^/2Cl^−^ symporter gene in M. jelskii initially increases ≈12-fold during the hyperosmotic challenge, transcription clearly underpinning Cl^−^ secretion. Gene expression of the Na^+^/K^+^/2Cl^−^ symporter in *Neohelice granulata* suggests that salt secretion also is associated with increased Na^+^/K^+^-ATPase activity during high salinity challenge (45 ‰S, Luquet et al., 2005). Thus, while Na^+^/K^+^-ATPase expression and activity are essential for both ionic hyper- and hypo-regulation, it is the insertion of newly translated, basally located Na^+^/K^+^/2Cl^−^ symporter molecules that likely drives the ability to secrete salt against a gradient in *M. jelskii*.

Interspecific comparison of gill ion transporter gene expressions shows that the basal V(H^+^)-ATPase expression in fresh water in *M. jelskii* is half that in *D. pagei*. Na^+^/K^+^-ATPase expression is 4-fold while Na^+^/K^+^/2Cl^−^ expression is similar. V(H^+^)- ATPase expression in *M. jelskii* increases only on hyper-osmotic challenge, suggesting a putative role in salt secretion. Increased overall Na^+^/K^+^-ATPase expression in *M. jelskii* seems crucial for coping with both hypo- and hyper-osmotic challenge while increased Na^+^/K^+^/2Cl^−^ expression appears to be particularly relevant to salt secretion during hyper-osmotic challenge.

These findings show that while mRNA expression of the same ion transporters appears to underlie the osmotic responses of two remotely related hololimnetic crustaceans from similar habitats, i. e., the crab *Dilocarcinus pagei* and shrimp *Macrobrachium jelskii*, their molecular and systemic regulatory mechanisms are not convergent and appear to have been driven independently. Each species has employed a distinct strategy during its lengthy evolutionary adaptation to fresh water, manifested in very different responses to hypo- and hyper-osmotic challenge at the gene transcription level and via consequent cellular and systemic adjustments.

## Acknowledgements

Crab and shrimp collections were authorized by the ICMBio/MMA (permit 29594-12 to JCM). We thank the managers at the Fazenda São Geraldo and Clube Náutico Araraquara for providing access to the crabs and shrimps, respectively. We are grateful to Drs Rogério Faleiros, Mariana Capparelli and Anieli Maraschi for assistance with fieldwork, and Susie Teixeira Keiko for technical assistance. We are indebted to Prof. José Antunes Rodrigues and Dr Lucila L. K. Elias (Departamento de Fisiologia, FMRP, USP) for laboratory support and use of the StepOnePlus Real- Time PCR Detection System. This investigation is part of a Ph D thesis submitted by MMM to the Graduate Program in Comparative Biology, Departamento de Biologia, FFCLRP, Universidade de São Paulo.

## Data availability

The datasets generated for this study are available on request to the corresponding author. Sequence data have been deposited with GenBank at the National Center for Biotechnology Information, USA.

